# Forecasting high pathogenicity avian influenza with a stochastic mechanistic model: performance and lessons for Australia

**DOI:** 10.64898/2026.08.24.746897

**Authors:** Meryl Theng, Simin Lee, Michelle Wille, Thao P. Le, Andrew C. Breed, Carlos Donoghue, Chris Baker, Simon M. Firestone

## Abstract

High pathogenicity avian influenza (HPAI) H5N1 clade 2.3.4.4b has caused a global panzootic with unprecedented impacts on wildlife and livestock, making evidence-based disease mitigation and outbreak response critical. In this paper, we describe a spatiotemporal mechanistic model of infectious disease dynamics developed for the HPAI Modelling Challenge and its implications for forecasting and policy in Australia. To emulate emergency response conditions, we adapted an existing model for rapid deployment rather than developing a bespoke model. We refined the model iteratively across the challenge to better analyse the provided outbreak data. Throughout the challenge, we accurately forecast temporal trends and local outbreak spread, but could not predict rarer, long-distance dispersal events. The challenge ended before HPAI H5N1 was first detected in Australia (June 2026), providing a critical opportunity to test our response modelling readiness for an incursion in wildlife and potential spillover into commercial poultry. Our experience identifies three key considerations for Australia’s HPAI H5N1 preparedness: targeted enhancements to our model to improve forecast precision and enable scenario-based policy evaluation; the critical value of pre-existing modelling infrastructure for rapid emergency response; and sustained collaboration between research and policy institutions to align modelling capabilities with outbreak response requirements.

## 1. Introduction

High pathogenicity avian influenza (HPAI) H5N1 clade 2.3.4.4b (hereafter HPAI) has caused a global panzootic, with unprecedented impacts on wildlife (Wille and Waldenström, 2023) and multiple livestock sectors (WOAH, 2026). Globally, hundreds of millions of poultry have died or been culled in response to infection with HPAI, and over 1000 dairy herds across 20 US states have been infected (WOAH, 2026; CDC, 2026; USDA APHIS, 2026). Development of evidence-based measures for disease mitigation and outbreak response and management is crucial to manage a disease event of this scale. Modelling has become an indispensable part of epidemiology, providing crucial insights into the understanding and management of complex pathogens such as HPAI (Wang et al., 2026). Specifically, modelling the epidemiology of HPAI may provide predictions on possible futures allowing for advanced warning, risk-based biosecurity preparedness, and critically, potential short- and long-term outcomes to management decisions (Gourram et al., 2023). In real terms, examples where modelling has been well integrated into HPAI responses in real-time are rare. Recent examples include forecasting exposure and spread among wild waterfowl in North America to provide advanced warning to poultry producers (McDuie et al., 2024), and predicting spatiotemporal spill-over risk from wild birds in the USA (Prosser et al., 2024). Modelling such as this identifies risk periods associated with the timing of bird migration and highlights when biosecurity should be enhanced.

At the time the challenge was undertaken (January-April 2026) (WiLiMan, 2026), Australia remained the only continent which was free from HPAI H5N1 (Wille et al., 2026b), though it has experienced multiple HPAI H7 outbreaks resulting from spilling over of low pathogenicity avian influenza (LPAI) H7 from local wild birds to commercial poultry with subsequent mutation to HPAI, albeit of a much smaller and more manageable scale (Breed et al., 2024; Wille et al., 2026a). Unlike other regions (e.g., North America and Europe), in Australia, integration of modelling activities for HPAI preparedness or response is relatively limited and is under active development. Critically, Australia lacks real-time forecasting models to guide HPAI outbreak response, a gap highlighted by the utility of such models during the COVID-19 pandemic (Moss et al., 2023, 2026). Whilst forecasting models for other livestock pathogens in the Australian landscape have recently been developed (e.g., equine influenza; Theng et al. 2024), re-purposing these models for HPAI required further work.

Building, testing, and refining models is time-consuming and rarely feasible during active outbreak response. Modelling challenges offer a valuable opportunity to develop and validate forecasting approaches during peacetime, strengthening preparedness before real epidemics emerge. Following previous challenges on Ebola, dengue, and African swine fever (Viboud et al., 2018; Johansson et al., 2019; Ezanno et al., 2022), the HPAI Modelling Challenge continued this tradition by providing a synthetic outbreak setting with realistic poultry production systems and control interventions (WiLiMan, 2026). This presented an ideal opportunity for us to adapt existing models and refine workflows for fitting them to an HPAI scenario— knowledge and tools we could later consolidate for Australian HPAI outbreak response.

Herein, we describe the efforts of our cross-sectoral team where we adapted an existing spatiotemporal mechanistic model—originally developed for foot-and-mouth disease—to forecast HPAI dynamics as part of the challenge. We used this model to provide forecasts across the three challenge phases. We begin by describing the model structure and key adaptations for HPAI, and then our fitting (parameter estimation) approach using Approximate Bayesian Computation with Sequential Monte Carlo (Section 2). We report forecasts of the number of cases and spatial infection risk at the upcoming weeks and assess forecasts at Phases 1 and 2 (Section 3). We conclude with a discussion of model strengths, limitations, and refinements for future iterations; including lessons learned regarding rapid modelling capacity and implications for Australian HPAI outbreak preparedness (Section 4).

## 2. Methods

### 2.1. HPAI Modelling Challenge

We adapted a pre-existing model specifically for the HPAI Modelling Challenge, and the methodology described here represents work conducted entirely within the 3-month window of the challenge. Broadly, the aims of the challenge itself were to use mathematical modelling to provide forecasts and policy recommendations during an outbreak of the fictional “Jolly Island”. Our team comprised of mathematical modellers, epidemiologists and those undertaking avian influenza surveillance in Australia from The University of Melbourne, as well as policy makers and epidemiologists from the Australian Department of Agriculture, Fisheries and Forestry (DAFF).

The challenge organisers provided data at three time-points, one month apart, that we used to forecast temporal trends of case counts, create spatial risk maps, estimate key transmission parameters and make policy recommendations. Six primary datasets were provided: (1) population data (farm ID, geographic coordinates, species, production type, capacity); (2) outbreak records (farm ID, date of suspicion, date of confirmation, culling dates and status); (3) preventative culling records (farm ID, cull status, dates); (4) farm movements (date, source farm, destination farm, volume); (5) farm activity logs (farm ID, batch entry and exit dates, volume); (6) geographic information and boundaries (e.g., counties, land use, high-risk zone where spillover from wild birds is more likely) (WiLiMan, 2026). Additionally, scanned mortality ledgers from three infected farms provided information about intra-farm disease dynamics. We focused our analysis on population data, outbreak and preventative culling records. Specific farm production types (e.g., chickens –broilers and layers, ducks – conventional and organic), movement and activity data were not used.

A key component of the scenario was the presence of two susceptible commercial poultry species: chickens and ducks. Therefore, we needed a model that has multiple species and allows for differences in transmissibility between species. We chose to adapt an existing foot-and-mouth disease (FMD) model (Lee et al., 2026b), because it already includes multiple species with different transmission characteristics. The model infers unobserved parameters by fitting to outbreak data, then uses these to make projections; thus, it does not carry across epidemiological parameters from FMD, only the general model structure. The primary adaptation for this challenge was configuring the model’s multi-species framework to represent chicken and ducks (see next section 2.2).

### 2.2. Model description

Our existing epidemic model is a stochastic model of inter-farm disease transmission, influenced by species-specific abundance of susceptible animals, distance to other premises, and intra-farm Susceptible-Exposed-Infectious-Recovered (SEIR) dynamics, with reactive culling implementation.

At time *t*, the total force of infection experienced by susceptible farms is:

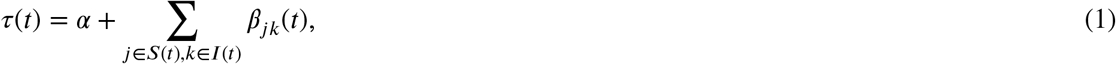

which is the sum of the individual infection pressures *β*_*jk*_(*t*) acting on all susceptible premises *j* from all infectious premises *k* at time *t* and *α* is the background transmission rate.

The infection pressure exerted by each pair of individual infectious and susceptible premises is

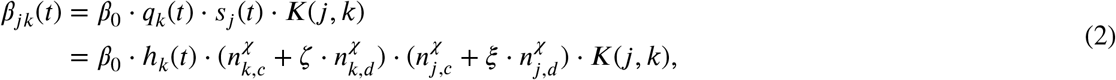

where *β*_0_ is the baseline transmission rate, ***K*** (*j,k*) is a spatial kernel describing the transmission rate decay with increasing distance, and *q*_*k*_(*t*) and *s*_*j*_ (*t*) denote the infectivity of farm *k* and the susceptibility of farm *j*, respectively. The latter two terms depend upon the total number of animals of each type on each farm and covariates/parameters *ζ* and *ξ*. Originally developed for FMD for cattle, pig, sheep, and others (Lee et al., 2026b), the key adaptation made for HPAI was to include transmission for chicken and duck species only. *n*_*k,c*_ and *n*_*k,d*_ denote the number of chickens and ducks, respectively, held by each premises *k* (or *j*). We assumed these values equal the farm capacity reported in the population data. *ζ* and *ξ* represent the relative infectiousness and susceptibility, respectively, of chickens farms to duck farms after accounting for the effect of numbers of animals on each farm with the term *χ* which allows for non-linear relationships. *h*_*k*_(*t*) is the prevalence of infectious individual animals within farm *k* at time *t* 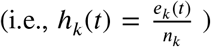, obtained from the solution of an intra-farm SEIR model (with intra-farm parameters *β*_intra_, *σ*_intra_, and *γ* _intra_ as the transition rates between states). Finally, we adopted a heavy-tailed, Cauchy-type transmission kernel, 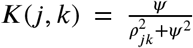, where 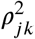 is the squared Euclidean distance between the centriods of farms *j* and *k*, and *Ψ* is a scaling parameter.

At time *t*, new infections—susceptible farms (*S*) transitioning to the exposed state (*E*)—are determined based on the following stochastic process:

1. The number of new infections proposed to occur is sampled from a Poisson distribution with mean *τ* (*t*) (Eq. 1), giving *E*(*t*) ^*^ ~Pois(*τ* (*t*)).
2. Proposed new infectees (*E*(*t*)^*^) are sampled from the set of premises susceptible at that time, *S*(*t*), probabilistically weighted based on the infection pressure being exerted on them by all the infectious premises at that time, *I*(*t*).
3. The proposed infectors (*I*(*t*)^*^) of these new infectees are sampled from *I*(*t*) probabilistically weighted based on the infection pressure that they are each exerting on each of the proposed infectees at time *t*.
4. The Poisson process is locally-thinned (Karr, 1991), where each new infection is only allowed to occur if the prevalence of infection on the proposed infector (*i* ∈ *I*(*t*)^*^) is greater than a random draw, u, from a uniform distribution *u* ~ *V* (0, 1), i.e., if *h*_*k*_(*t*) *> u*, then infection occurs at *j* ∈ *E*(*t*)^*^.

Farms in the exposed state (*E*) transition to being infectious (*I*) after the latent period, which was assumed to follow a Gamma distribution reflecting a mode of one day and maximum of two days (Lambert et al., 2023). Both these states are unobserved events, so we assumed the incubation period—time between exposure to first clinical signs observed (*t*_*k*,clinical_ – *t*_*k,e*_)—followed a Gamma distribution with a mode of 2.5 days and maximum of 10 days (Lambert et al., 2023). We note that these periods are species-specific (i.e., ducks have longer latent and presymptomic periods than chicken) but used the same assumption generalised across all chicken and duck farms.

The farm remains infectious (*I*) until it is depopulated (D) or recovered (R), whichever occurs first. The time to depopulation after notification (t_*k*_,depop – t_*k*_,notified) is sampled from a uniform distribution *V*(0, *dtND*_max_), where *dtND*_max_ is the maximum observed delay from notification to depopulation.

All notified farms are flagged for depopulation upon notification (i.e., representing reactive culling). An infectious farm can recover when the number of infected animals within it is modelled to be *<* 0.5 at time *t*, obtained from the solution of the intra-farm SEIR model.

Reporting delays—the time to notification after first clinical signs (*t*_*k*_,notified – *t*_*k*_,clinical)—were similarly drawn from a uniform distribution *V*(0, *dtCN*_max_), where *dtCN* _max_ is the maximum observed delay from clinical signs to notification.

We incorporated preventative culling as observed in the data provided. Thus, susceptible farms selected for culling and their culling dates were identical across all simulations.

### 2.3. Parameter estimation

The model parameters were either estimated from the outbreak data, taken from a review of avian influenza models (Lambert et al., 2023), or estimated from the model fitting process (Table 1). Naïve bounds for the flat priors placed on the inter-farm parameters, were first taken directly from the FMD model and then refined based on preliminary model runs on the HPAI challenge data.

**Table 1.**
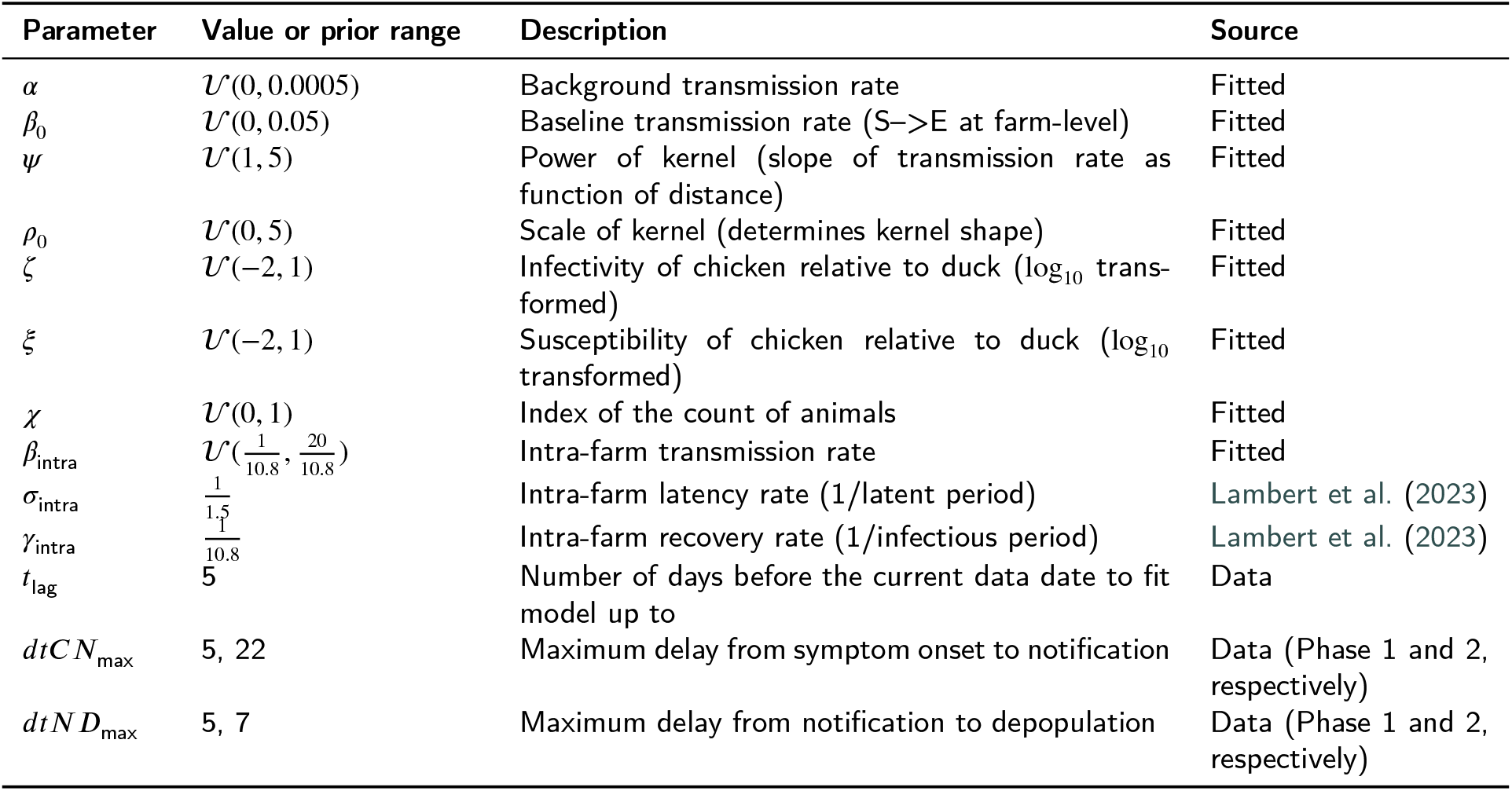
Parameters used in the model and their values/ranges.

| Parameter | Value or prior range | Description | Source |
| --- | --- | --- | --- |
| $\alpha$ | $\mathcal{U}(0, 0.0005)$ | Background transmission rate | Fitted |
| $\beta_0$ | $\mathcal{U}(0, 0.05)$ | Baseline transmission rate (S→E at farm-level) | Fitted |
| $\psi$ | $\mathcal{U}(1, 5)$ | Power of kernel (slope of transmission rate as function of distance) | Fitted |
| $\rho_0$ | $\mathcal{U}(0, 5)$ | Scale of kernel (determines kernel shape) | Fitted |
| $\zeta$ | $\mathcal{U}(-2, 1)$ | Infectivity of chicken relative to duck ($\log_{10}$ transformed) | Fitted |
| $\xi$ | $\mathcal{U}(-2, 1)$ | Susceptibility of chicken relative to duck ($\log_{10}$ transformed) | Fitted |
| $\chi$ | $\mathcal{U}(0, 1)$ | Index of the count of animals | Fitted |
| $\beta_{\text{intra}}$ | $\mathcal{U}(\frac{1}{10.8}, \frac{20}{10.8})$ | Intra-farm transmission rate | Fitted |
| $\sigma_{\text{intra}}$ | $\frac{1}{1.5}$ | Intra-farm latency rate (1/latent period) | Lambert et al. (2023) |
| $\gamma_{\text{intra}}$ | $\frac{1}{10.8}$ | Intra-farm recovery rate (1/infectious period) | Lambert et al. (2023) |
| $t_{\text{lag}}$ | 5 | Number of days before the current data date to fit model up to | Data |
| $dtCN_{\text{max}}$ | 5, 22 | Maximum delay from symptom onset to notification | Data (Phase 1 and 2, respectively) |
| $dtND_{\text{max}}$ | 5, 7 | Maximum delay from notification to depopulation | Data (Phase 1 and 2, respectively) |

We used Approximate Bayesian Computation with Sequential Monte Carlo (ABC-SMC) to approximate posterior distributions of estimated parameters (Beaumont et al., 2009). The algorithm samples candidate parameter sets from prior distributions, simulates model outputs, and evaluates the ‘closeness’ between these and the observed data using distance functions, in this case: (i) temporal distance – sum of absolute differences in daily case counts (i.e., newly exposed farms); and (ii) spatial distance – correlation between grid-cell case counts. Parameter sets satisfying distance thresholds are accepted as particles. The SMC algorithm improves efficiency by progressively decreasing tolerance thresholds across sequential generations, narrowing the parameter space explored (Beaumont et al., 2009). Accepted particles form the posterior distribution and model fit, while forward projections from these particles generate forecasts. This approach is detailed in Theng et al. (2024).

Simulations commenced on 17 December 2025, initialised with confirmed infections on farms on Jolly Island, prior to the first case notification (22 December 2025). The model was fitted to case data from the first notification to several days before each phase cutoff to account for notification delays, which were typically within 3-5 days of clinical signs (Fig. S1 in Supplementary Material). For Phase 1, the fitting period spanned 22 December 2025 to 11 January 2026 (3 days before cutoff). For Phase 2, the fitting period initially spanned 22 December 2025 to 9 February 2026 (5 days before cutoff), then subsequently refitted from 30 January to 9 February (see section 3.1). We targeted 1,000 accepted particles per SMC generation across six generations. Random seeds were uniquely assigned to 50 cores on the University of Melbourne’s High-Performance Computing Cluster ‘Spartan’ (on the ‘sapphire’ partition, with Intel® Xeon® Gold 6448H processor) and recorded for reproducibility. Model outputs comprised temporal forecasts of daily case trajectories, spatial forecasts of two-week-ahead farm-level infection risk at specified time-points and marginal distributions for inferred parameters from the estimated posterior.

## 3. Results

Our model converged at SMC generation 5 for Phase 1, yielding 738 accepted particles. For Phase 2, after addressing fitting challenges (see below), convergence was achieved at generation 4 with 1,000 accepted particles. These particles formed the basis for all submitted forecasts and analyses. This required approximately 48 hours of runtime per phase (including job queue wait times); however, particles from earlier generations (completed in substantially less time) could have provided meaningful predictions under tighter time constraints, demonstrating the flexibility of the ABC-SMC approach for rapid response scenarios (see Figs. S2, S3 in Supplementary Material).

### 3.1. Overcoming model fitting challenges in Phase 2

During Phase 2, the model fitted with Phase 1 parameters (described in section 2.3) yielded a poor fit with the observed data, undermining forecast reliability. We systematically explored several refinements: (i) stricter temporal distance criteria (e.g., absolute differences in daily cases must not exceed the threshold), (ii) an additional distance function constraining peak timing, and (iii) adjustments to disease parameters (latent, incubation period) and prior bounds. These approaches made negligible difference to model fit. We ultimately resolved this by adjusting the fitting window: initialising simulations with data prior to 30 January 2026 (rather than 22 December 2025) and fitting to data up to five days before the current date. By narrowing the fitting window to a period with more consistent control measures, we avoided fitting across multiple intervention changes, substantially improving model-data fit (Fig. S4 in Supplementary Material).

### 3.2. Forecast performance (Phase 1 and 2)

Forecasts at both Phase 1 and Phase 2 captured the observed daily case trajectories within the 95% prediction intervals (Fig. 1). Delaying the time of forecast by three and five days proved sumcient to account for reporting delays in Phase 1 and 2, respectively. **Phase 1 forecasts** showed substantial variability across simulations, with most trajectories continuing to grow and peak beyond the current date, whilst few died out completely. The realised outbreak tracked towards the lower bounds of the prediction intervals, particularly at longer lead times. **Phase 2 forecasts** were substantially more accurate. Simulations predicted outbreak resolution between 31 January and 19 April 2026 (95% prediction interval), with the observed epidemic trajectory falling within these bounds.

**Figure 1.**
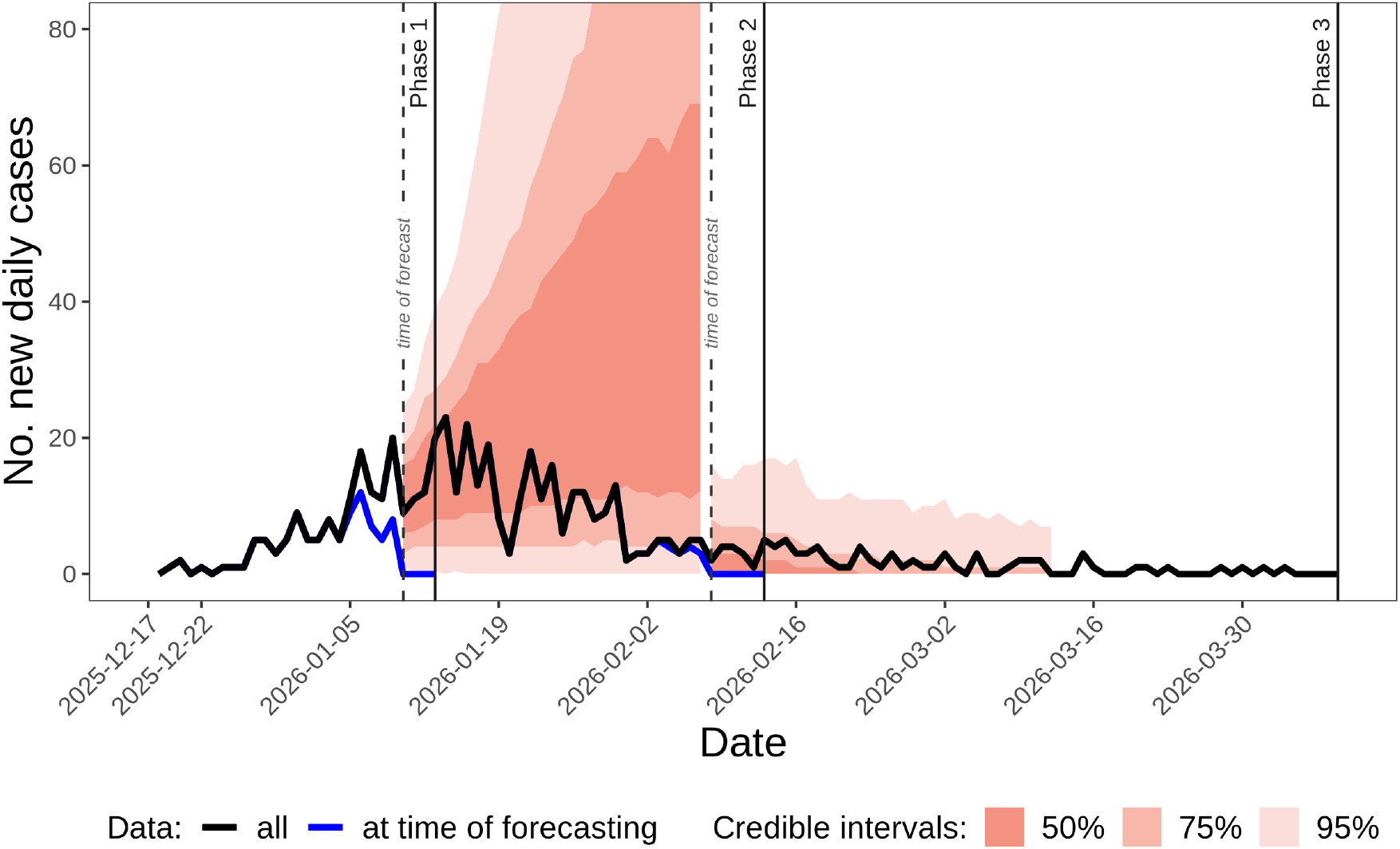
Forecasts of daily case incidence (newly infected premises) at Phases 1 and 2. Red bands show prediction intervals (50%, 75%, and 95%) from forecasts made three and five days before Phase 1 and 2 cutoff, respectively, to account for reporting delays. The black line represents all observed case counts (ground truth), whilst blue lines show the reported cases available at each forecasting time (the data used to fit the model).

Spatial forecasts were able to capture the extent of local outbreak spread, but not longer-distance dispersal events—noting that these were not explicitly modelled (Fig. 2). The **Phase 1** forecast was able to capture continued spread in the more immediate vicinity of the southern outbreak cluster, including some continued spread in the north (Fig. 2a). **Phase 2** predictions captured observed cases near the southern extent of the northern cluster (Fig. 2b).

**Figure 2.**
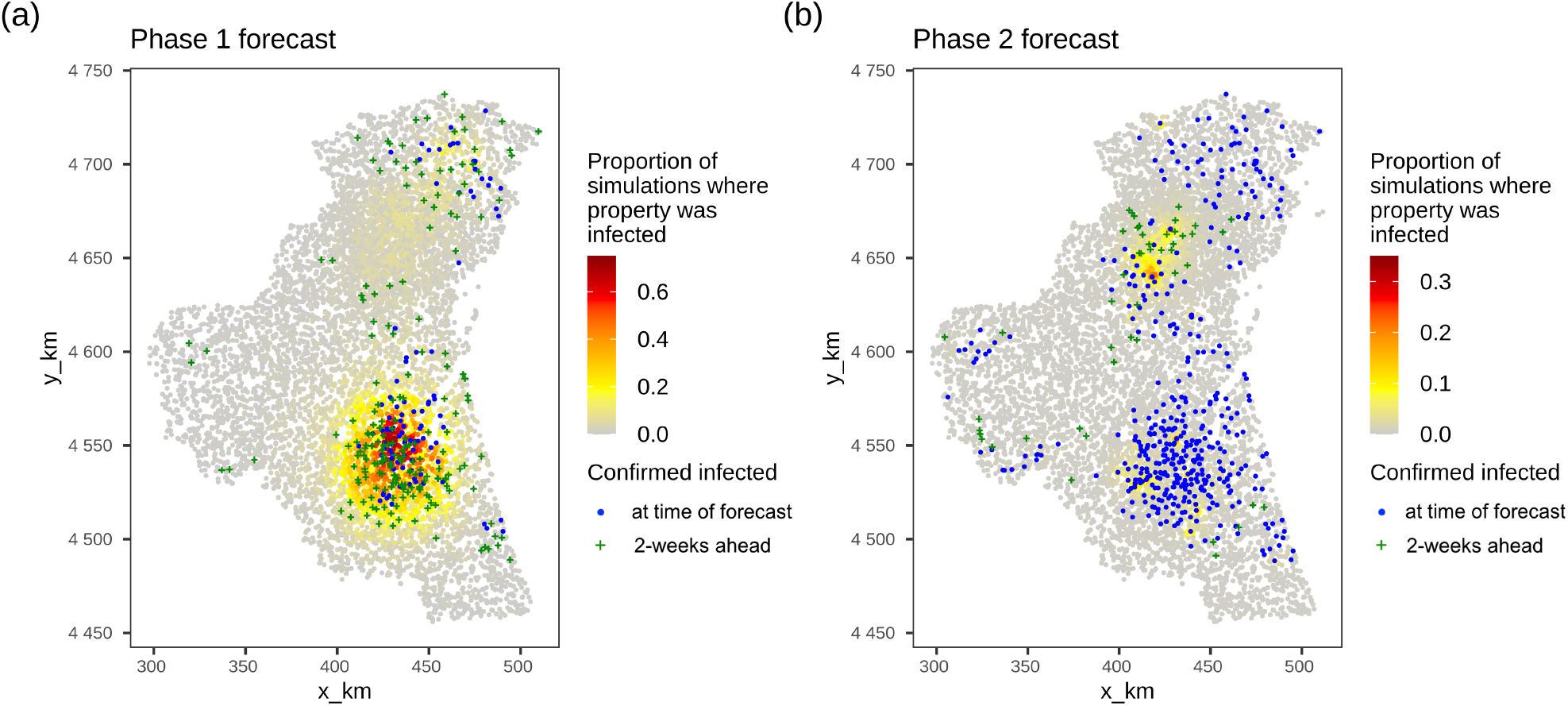
Forecasts of farm-level infection risk in the two-weeks ahead for Phases 1 and 2. Maps show the proportion of simulations (out of 738 and 1000 possible futures respectively) in which a farm (each point on the map) is infected between the simulation seeding date (phase 1: 2025-12-22; phase 2: 2026-01-30) and the *2-weeks ahead* of the end of each phase. Blue points represent premises that were infected (as of the time of forecasting) and green crosses are farms that were infected during the forecasting period.

### 3.3. Species contribution to infections (Phase 1)

In Phase 1, we were asked to characterise the relative contribution of the chicken farms to virus spread as compared to duck farms.

Posterior estimates of ζ and ξ (infectivity and susceptibility of chicken relative to duck premises, respectively; Eq. 2) indicated duck farms were 2.3-fold [0.21–33] more infectious and 5.9-fold [0.15–100] more susceptible than chicken farms (Fig. 3) on a per capita basis (i.e., assuming chicken and duck populations are of the same size). Though for ξ, the limited divergence from the uniform prior suggests this effect is weak. χ (Eq. 2) was low, reflecting the large numbers of animals of each species on most farms and its non-linear effect on farm-level infectivity and susceptibility.

**Figure 3.**
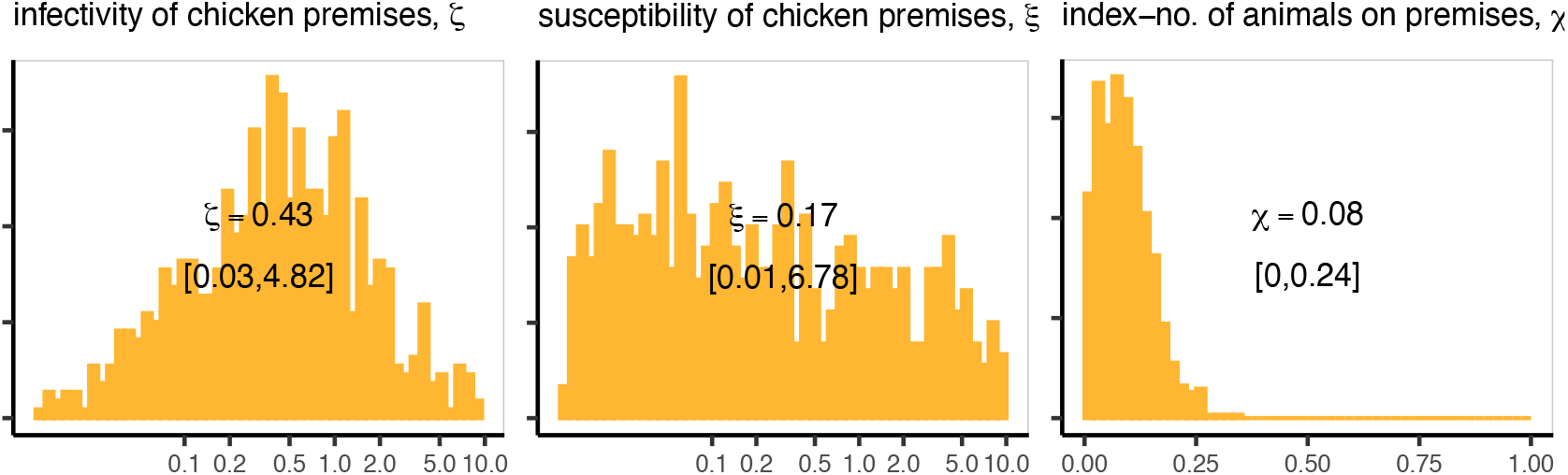
Posterior distributions of parameters relating farm abundance to transmission dynamics, assuming farms operate at full capacity throughout. Median estimates with 95% credible intervals (in square brackets) are shown for each parameter.

### 3.4. Counterfactual analysis (Phase 3)

In Phase 3, rather than forecasting future epidemic trajectories, we were asked to address a counterfactual scenario, where we estimated the number of outbreaks averted by the preventive culling strategy.

We fitted the model with data up to 5 January 2026 (the first preventive culling event) and generated counterfactual forecasts assuming no preventive culling intervention. Comparing predicted trajectories to the observed outbreak revealed a substantially larger epidemic would have occurred without this strategy (Fig. 4). We estimate that 881–8192 infections (95% CI) were averted in total. This should be interpreted as an upper bound, as the counterfactual also reflects the combined effects of all concurrent control measures (e.g., reactive culling, zoning, movement restrictions), which were not explicitly modelled.

**Figure 4.**
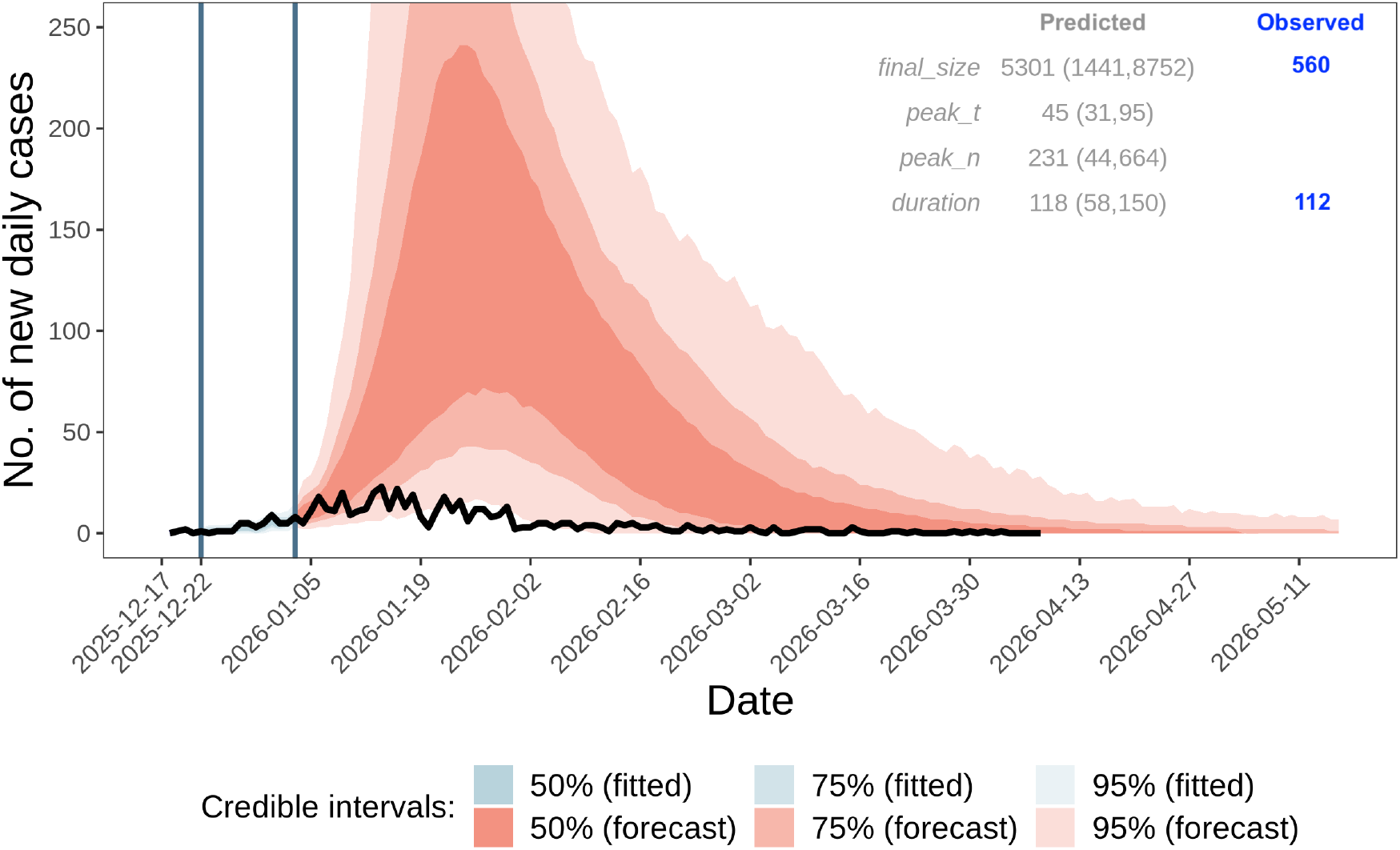
Forecast of daily newly exposed properties under a counterfactual no-intervention scenario. Shaded bands show the 50%, 75%, and 95% prediction intervals from 1000 possible futures simulated by the model. Blue shading denotes the model fit to observed data, while red shading represents the counterfactual forecast assuming no preventative culling or other control measures. The black line shows the observed epidemic trajectory (Phase 3), reflecting the implementation of interventions. Vertical blue lines indicate the fitting window, from first detection (2026-12-22) to the day before the first preventative culling event (2026-01-05); earlier data (black line) were used for model initialisation. Values in the top-right corner summarise predicted (median and 95% CI) and observed metrics: final outbreak size (final size), peak timing (days), peak incidence, and outbreak duration (days).

## 4. Discussion

We adapted an existing stochastic mechanistic modelling framework to forecast the simulated outbreak in the HPAI Modelling Challenge. Our model successfully captured temporal trends in case incidence and the spatial extent of local outbreak spread, though it was not setup to predict long-distance dispersal events. Here, we discuss the model’s strengths and limitations, refinements planned in future research and implications for Australian HPAI H5N1 preparedness.

### 4.1 Strengths and limitations, refinements for the future

Our approach demonstrated several key advantages for rapid response modelling. First, rapid adaptability of an existing modelling framework, possessing a multi-species structure and reactive culling framework closely aligned with HPAI requirements. Critically, the ABC-SMC fitting algorithm required no modification—a likelihood-free Bayesian approach that avoids the time-consuming reprogramming of training algorithms necessary for full MCMC approaches (Drovandi et al., 2016). This enabled deployment of a functional forecasting tool within two weeks or five person-days of development effort. Second, Bayesian inference of unknown parameters eliminated the need for manual calibration, allowing the model to learn directly from outbreak data. This is particularly valuable as transmission parameters are often uncertain and context-specific, shaped by disease characteristics, population structure, and evolving behaviours and interventions (Scarpino and Petri, 2019). Third, operational feasibility; whilst results presented here required ~ 48 hours to prepare per phase, earlier SMC generations provided meaningful predictions after only 4-10 hours of analysis (Figs. S3, S4 in Supplementary Material), demonstrating flexibility for meeting the tighter time constraints expected in genuine large-scale outbreak responses.

We acknowledge several key limitations of our approach that constrained forecast accuracy. First, omission of production activity and inter-farm movements likely reduced temporal and spatial forecast precision. We assumed farm abundance equalled full capacity, so farms with periods of zero activity continued to contribute to disease transmission throughout the outbreak. We also assumed a homogeneous farm network structure, preventing prediction of long-range dispersal events seeding new outbreak clusters. This is a critical gap to fill, as movements represent a primary transmission pathway, particularly for stage 1 to stage 2 chicken broiler farm connections (WiLiMan, 2026). Second, preventive culling was not explicitly modelled. Although we incorporated preventive cull data into simulations, the selection process (which farms to cull) was not mechanistically represented as this would have taken considerable time to code and verify for our first use in this context. Consequently, Phase 1 forecasts tended to overestimate outbreak size, likely representing what could have happened without this intervention measure and other measures not modelled (e.g., movement restrictions and enhanced biosecurity). Third, species-specific parameters did not reflect observed patterns. Despite fitting parameters for chicken infectivity (*ζ*) and susceptibility (*ξ*) relative to ducks, posteriors suggested duck farms were far more infective and more susceptible than chicken farms, contradicting the observed outbreak where chicken farms dominated confirmed case numbers. This coupled with the weak divergence of *ξ* from its naïve prior suggests the model struggled to learn species-specific differences, likely because our fitting criteria did not discriminate between farm types.

From this experience, we identify several priority developments to enhance model performance before application in a real HPAI outbreak in Australia:

- **Explicit representation of preventative culling:** Mechanistically representing on-the-ground culling decisions (e.g., preventative culling of all farms within 3 km of newly confirmed cases) would allow forward projections to incorporate the strategy, reducing the tendency to overestimate outbreak growth in the exponential phase, and also enable genuine scenario-based policy evaluation (e.g., Probert et al. 2018; Howerton et al. 2023). To date, Australia has never leveraged a preventative culling strategy against HPAI H7 outbreaks.
- **Modelling of movements and farm activity:** Movements can be modelled via conditional movements based on farm ownership, farm type and production cycle (e.g., movements from chicken stage 1 broilers to stage 2 broilers to slaughterhouses), strength of connections based on contact history (Jewell et al., 2009), or allowing incorporation of real-time movement data (e.g., Paploski et al. 2021; Yoo et al. 2021).
- **Time-dependent background or baseline transmission rates:** Modelling these rates as time-varying (e.g., Funk et al. 2018), such that the model is flexible enough to accommodate changes in transmission rates that might occur due to unobserved processes. This may have improved the Phase 2 model-data fit to the longer period where the outbreak situation was continuously changing due to multiple interventions. This is particularly useful in real outbreak situations where it is practically impossible to fully observe and much less model the combination of behavioural changes and intervention in real-time (Scarpino and Petri, 2019).
- **Species-specific fitting criteria:** Incorporating separate ABC-SMC distance metrics for chicken and duck farms would allow the model to learn differences in transmission dynamics—e.g., fitting to chicken and duck case counts independently could improve inference of *ζ* and *ξ*. Also, different model formulations could be tested that focus on infectivity and susceptibility at the farm-level rather than fitting parameters at the animal level then accounting for farm size. This would be particularly useful in the Australian context given the implication of emu farms in the HPAI H7 outbreaks in 2020.
- **Spatially-varying spillover rates:** The high-risk zone (HRZ) for wild bird spillover (WiLiMan, 2026) should have a distinct background spillover rate (a; Eq. (1)), potentially evolving into a raster-based approach varying across the landscape (e.g., based on wild bird density, water body proximity, or other ecological covariates).

The first three developments require substantial development of the core simulation model, while the fourth should be straightforward to implement within the ABC-SMC algorithm, and the fifth can leverage an existing prototype for lumpy skin disease in cattle (LSD) that incorporates vector distribution (Lee et al., 2026a).

### 4.2. Learnings for Australia

Australia, the last continent to be infected with HPAI H5N1, confirmed its first detections of the virus in June 2026, predominantly in Southern Ocean seabirds along the southern coastline (DAFF, 2026a). Prior to the arrival of the virus to Australia, the Australian Government in collaboration with non-government agencies have undertaken considerable preparedness work through the formation of an “HPAI Taskforce” in 2024, which have undertaken simulation exercises such as Exercise Volare (DAFF, 2024), performed evaluations for surveillance systems in wildlife and commercial poultry, strengthened vaccination policy, and enforced strict national border controls (DAFF, 2026b).

Herein, we leveraged a livestock-to-livestock transmission model, which is designed to be generalisable across poultry systems. The modelling framework used here can be applied directly to the Australian poultry network given appropriate population data; notwithstanding key structural diGerences such as sparser premises network and a more centralised, vertically integrated supply chain relative to North America or Europe (Scott et al., 2009). Explicitly incorporating contact structure beyond spatial proximity between farms may nonetheless require tailoring to the Australian context. Finally, Australia’s poultry population includes ratites (notably emus) rather than the duck farms represented in the model; the duck farm abundance terms (*n*_*k,d*_ and *n*_*j,d*_ in Eq. 2) can readily represent these alternative species.

Irregular patterns of wild bird movement will largely determine when and where spillover from wildlife into Australian poultry occurs. Australia is inherently diGerent from the northern hemisphere in ways that are critical for modelling HPAI spillover, so spillover risk models developed overseas (e.g., Prosser et al. 2024; Liu et al. 2025) are not fit-for-purpose here. Unlike in the northern hemisphere, waterfowl—the main dispersers of low pathogenicity avian influenza (LPAI) and HPAI—do not follow predictable seasonal migration patterns, but are instead nomadic, tracking irregular rainfall around the Australian landscape (Ferenczi et al., 2021). Consequently, LPAI prevalence in Australian waterfowl varies strongly between years: in wet years, high recruitment of immunologically naïve juveniles drives high prevalence, whereas in dry years reproduction is reduced such that there is little or no recruitment of immunologically naïve birds (Ferenczi et al., 2016; Wille et al., 2023). These dynamics also influence the timing of LPAI and HPAI outbreaks in poultry (Ferenczi et al., 2021; Wille et al., 2026a). The pronounced seasonal spillover risk profile observed in Europe and North America is therefore absent, or only weakly expressed, in Australia. Consequently, spillover risk models for Australia must explicitly account for its distinctive avian influenza ecology.

This challenge provided a critical opportunity to adapt, implement and validate our existing livestock transmission model on a simulated HPAI outbreak. The learnings from this challenge are instrumental for Australia’s preparedness. First, we have identified specific model enhancements to improve our framework’s applicability to HPAI H5N1, and to address the types of questions posed by policy makers during outbreak response (detailed in section 4.1). Second, we demonstrated how prior investment in reusable modelling infrastructure can accelerate preparedness and response. Within approximately five person-days, our team had modified the existing FMD framework to represent HPAI transmission dynamics, and produced the initial set of outbreak forecasts. In a real outbreak, the ability to adapt existing tools may be just as important as the models themselves, allowing modellers to focus on disease-specific questions rather than rebuilding core analytical systems. Given Australia’s distinct ecology and livestock production systems, Australia-tailored models are essential; we cannot simply extrapolate findings from overseas without accounting for local conditions. Finally, the challenge catalysed meaningful collaboration between multiple university teams and DAFF. Critically, by embedding policy makers within our team, we demonstrated how modelling could directly address outbreak management questions, positioning modelling as a valued component of response decision-making. Further, this partnership has established a foundation for sustained engagement in animal health emergency preparedness, enabling rapid mobilisation of modelling expertise when needed and fostering alignment between research capabilities and policy requirements.

## Acknowledgements

This research was supported by the *Enhancing Models for Rapid Decision-Support in Emergency Animal Disease Outbreaks* (HASTE) project, which received co-investment (doi.org/10.47486/DC110) from the Australian Research Data Commons (ARDC). The ARDC is enabled by the National Collaborative Research Infrastructure Strategy (NCRIS). This research was supported by The University of Melbourne’s Research Computing Services and the Petascale Campus Initiative. Michelle Wille is supported by an Australian Research Council Future Fellowship. The Melbourne WHO Collaborating Centre for Reference and Research on Influenza is supported by the Australian Government Department of Health, Disability and Ageing. We thank Martin Cyster for discussions about modelling vaccinations.

## CRediT authorship contribution statement

**Meryl Theng:** Conceptualisation, Methodology, Software, Formal Analysis, Writing – Original Draft Preparation. **Simin Lee:** Methodology, Software, Formal Analysis. **Michelle Wille:** Validation, Writing – Original Draft Preparation. **Thao P. Le:** Conceptualisation, Writing – Review & Editing. **Andrew C. Breed:** Supervision, Validation, Writing – Review & Editing. **Carlos Donoghue:** Validation. **Chris Baker:** Conceptualisation, Supervision, Writing – Original Draft Preparation. **Simon M. Firestone:** Conceptualisation, Methodology, Supervision, Writing – Review & Editing.

