## Supplementary Material for "Forecasting high pathogenicity avian influenza with a stochastic mechanistic model: performance and lessons for Australia"

### A. Supplementary

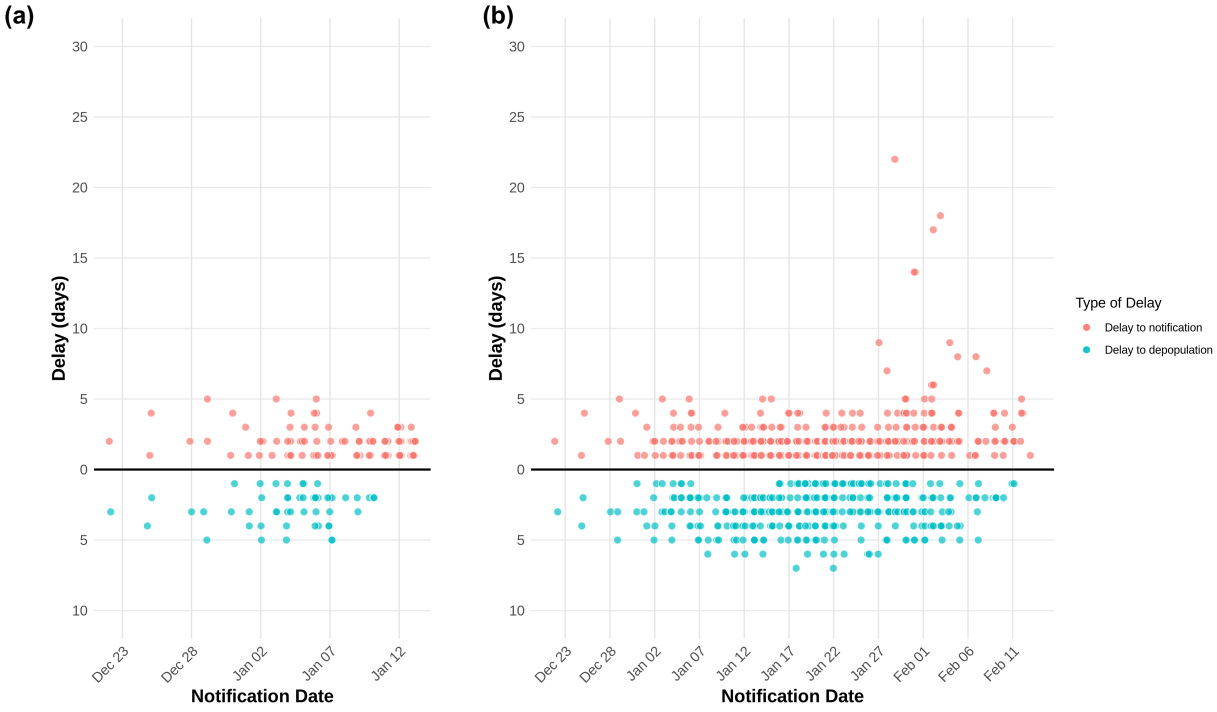


**Figure S1:** Observed individual farm delays by notification date from (a) phase 1, and (b) phase 2 data. Each point represents one farm (jittered for visibility).

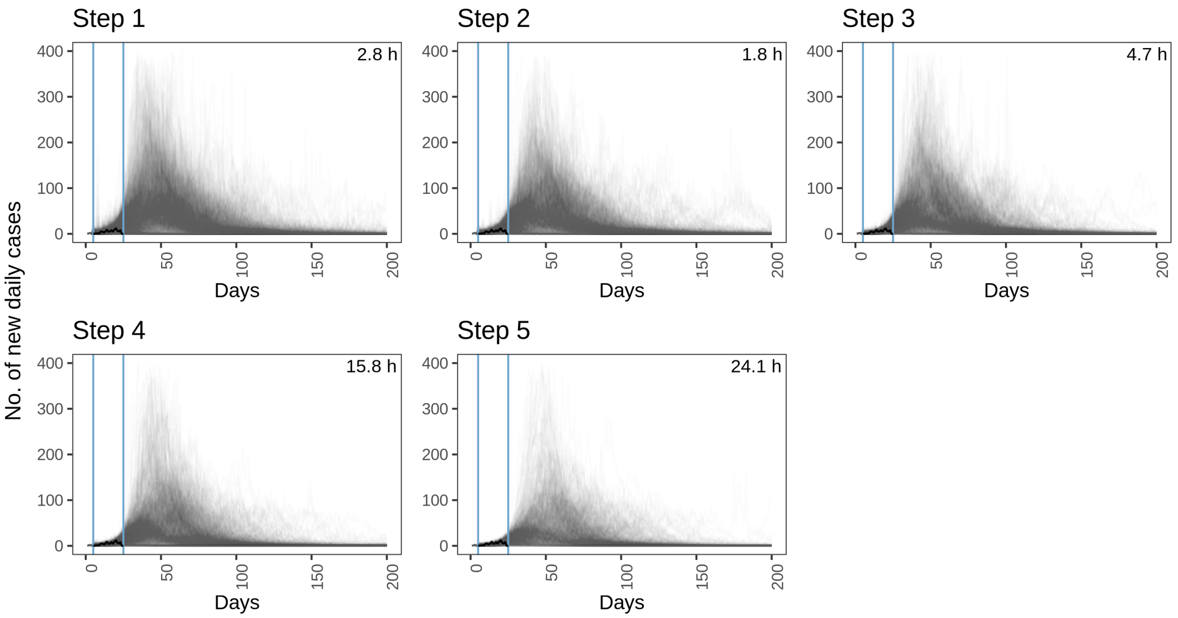


**Figure S2:** ABC-SMC step (generation) plots for **Phase 1**. Panels show daily case count trajectories (grey lines) simulated by accepted particles at each step, together with the time taken (in hours) to complete each generation (including job queue wait times). The black trend line is the observed data available at phase 1, and blue vertical lines indicate the fitting window. No particles were produced in step 6, which had a 24-h runtime limit.

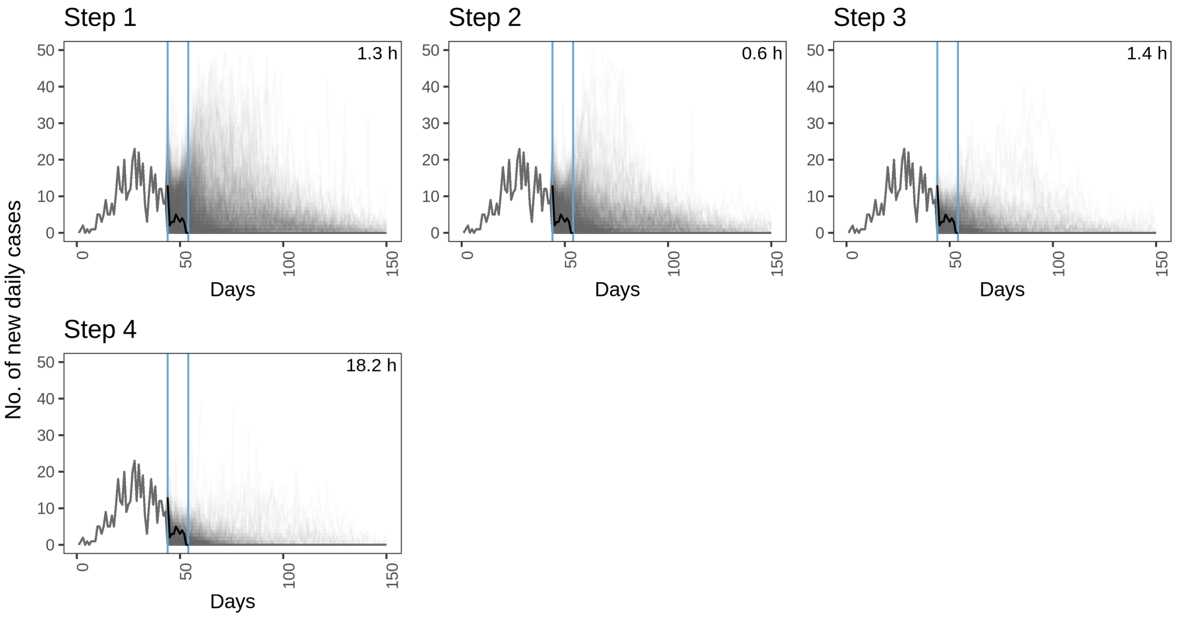


**Figure S3:** ABC-SMC step (generation) plots for **Phase 2**. Panels show daily case count trajectories (grey lines) simulated by accepted particles at each step, together with the time taken (in hours) to complete each generation (including job queue wait times). The black trend line is the observed data available at phase 1, and blue vertical lines indicate the fitting window. No particles were produced in step 5, which had a 24-h runtime limit.

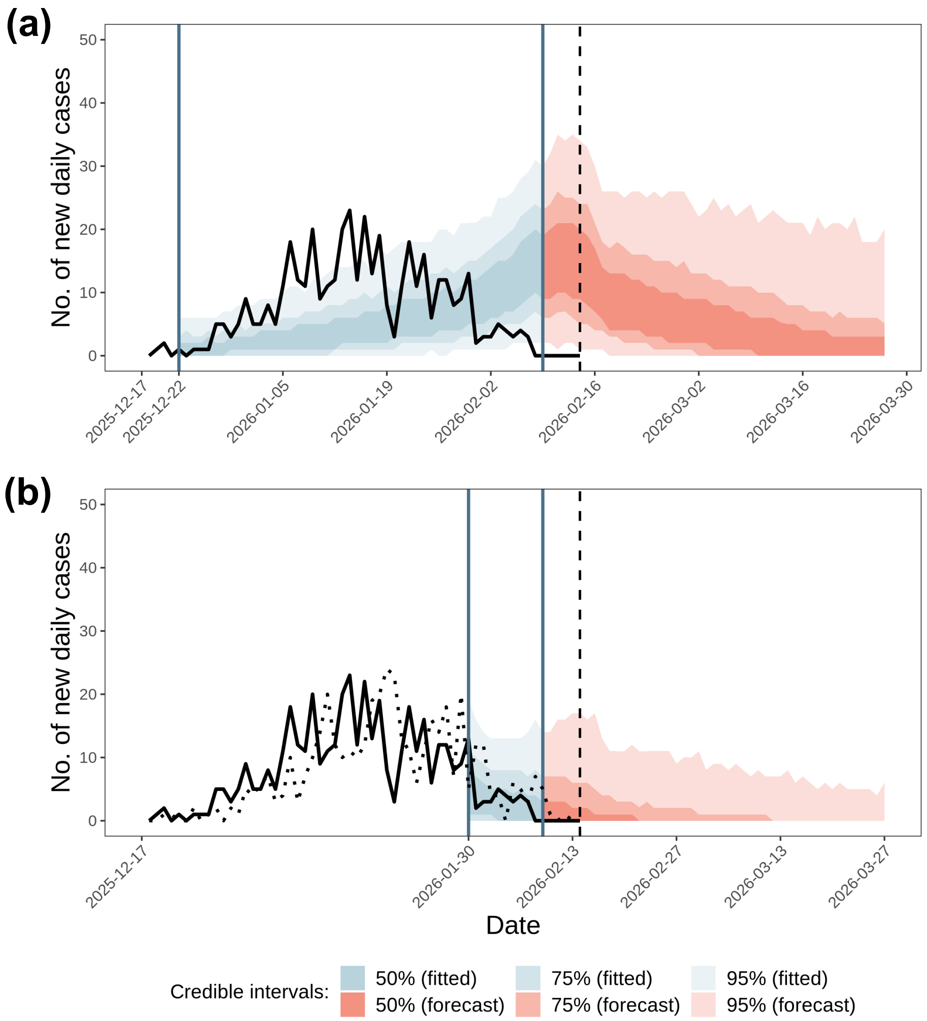


**Figure S4:** Fitted epidemic curves and forward projections (forecasts) from phase 2 data: (a) original fit to the data between the first notification date (22 December 2025) and five days before the phase cutoff (14 February 2026), and (b) improved fit to the data between 30 January 2026 and five days before the phase cutoff. The black line represents the number of new daily infections (i.e., new exposures) inferred from the observed notified cases (dotted line in b) based on the assumption that the incubation period is has a mode and maximum of one and two days, respectively.
